# Resolving systematic errors in widely-used enhancer activity assays in human cells enables genome-wide functional enhancer characterization

**DOI:** 10.1101/164590

**Authors:** Felix Muerdter, Łukasz M. Boryń, Ashley R. Woodfin, Christoph Neumayr, Martina Rath, Muhammad A. Zabidi, Michaela Pagani, Vanja Haberle, Tomáš Kazmar, Rui R. Catarino, Katharina Schernhuber, Cosmas D. Arnold, Alexander Stark

## Abstract

The identification of transcriptional enhancers in the human genome is a prime goal in biology. Enhancers are typically predicted via chromatin marks, yet their function is primarily assessed with plasmid-based reporter assays. Here, we show that two previous observations relating to plasmid-transfection into human cells render such assays unreliable: (1) the function of the bacterial plasmid origin-of-replication (ORI) as a conflicting core-promoter and (2) the activation of a type I interferon (IFN-I) response. These problems cause strongly confounding false-positives and -negatives in luciferase assays and genome-wide STARR-seq screens. We overcome both problems by directly employing the ORI as a core-promoter and by inhibiting two kinases central to IFN-I induction. This corrects luciferase assays and enables genome-wide STARR-seq screens in human cells. Comprehensive enhancer activity profiles in HeLa-S3 cells uncover strong enhancers, IFN-I-induced enhancers, and enhancers endogenously silenced at the chromatin level. Our findings apply to all episomal enhancer activity assays in mammalian cells, and are key to the characterization of human enhancers.

## The ORI is an inducible core-promoter in human cells

### Common plasmid DNA elements confound enhancer activity assays

While promoters are located at the 5’end of genes and initiate transcription locally, enhancers can activate transcription from distal core-promoters^1,2^. This defining property is frequently assessed in enhancer activity assays that test candidate DNA fragments outside their endogenous genomic contexts, which directly measures the candidate sequences’ enhancer functionality without the influence of the different flanking genomic regions. On a typical reporter plasmid for enhancer-activity assays (e.g. the pGL3/4 enhancer vectors), the candidate enhancer is placed downstream of a reporter gene or a barcode sequence (Fig. 1A), ensuring the assessment of bona fide enhancer- rather than promoter activity. Importantly however, in human cells luciferase reporter transcripts from the widely used pGL3/4 reporter system initiate predominantly in the bacterial plasmid origin-of-replication (ORI) rather than the minimal core-promoter^3^ (mCP, Fig. 1A). While the function of the ORI as a core-promoter is not unexpected given the presence of several core-promoter elements^3^ (Fig. S1A) and the ORIc’s propensity to remain nucleosome free^4^, it will likely impact enhancer-activity measurements: the undefined 5’ UTR resulting from uncontrollable and presumably cell type-specific splicing of the intervening~2 kb plasmid sequence will confound luciferase assay readouts. Moreover, differences in reporter transcript stability or transcriptional interference between the two core-promoters can also affect assays that measure reporter abundance at the RNA level, as all sequencing-based massively parallel reporter assays (MPRAs) including STARR-seq do^2,5,6^.

**Figure 1.**
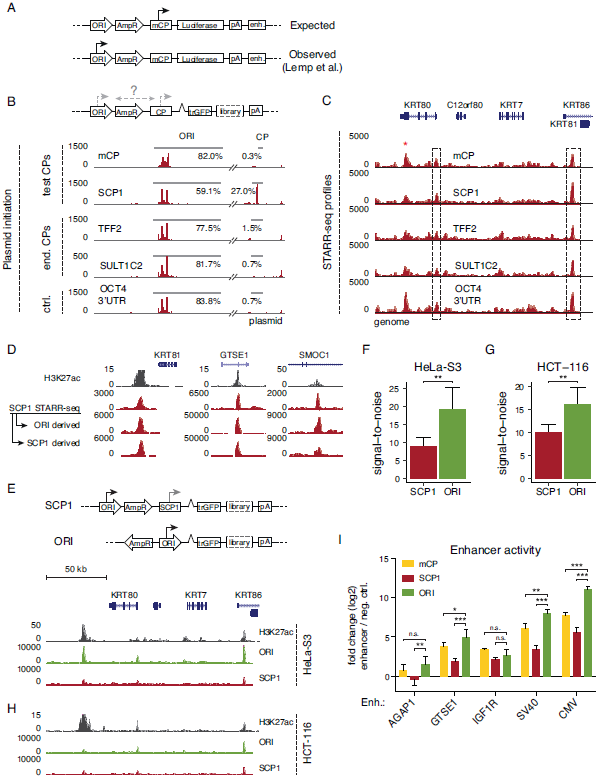
The ORI is an optimal core-promoter for STARR-seq and luciferase assays **A**, Typical layout of a reporter plasmid for enhancer activity assays (e.g. pGL3/4) with the origin-of-replication (ORI), a resistance gene (e.g. AmpR), a minimal core-promoter (mCP), a reporter gene (e.g. Luciferase), a polyadenylation sequence (polyA) and an enhancer candidate (enh.). The major site of reporter transcript initiation is indicated with an arrow (expected vs. observed according to ref. 3). **B**, Reporter transcript initiation on STARR-seq reporter plasmids as measured by STAP-seq for setups with two synthetic core-promoters (mCP, SCP1) and two endogenous core-promoters (TTF2, SULT1C2) vs. a negative control (ctrl., OCT4 3’UTR). Red vertical lines indicate transcription initiation sites with the respective initiation frequencies according to STAP-seq. The percentages indicate the fraction of all initiation events in either the ORI or the respective core-promoter. **C**, Representative STARR-seq enhancer activity profiles obtained for reporter setups from (B) are shown. The Refseq (GRCh37) gene track is indicated above, the dashed boxes indicate luciferase-validated enhancers. **D**, Representative enhancer activity profiles over three gene loci (indicated above; H3K27ac data from ENCODE, Table S2) for an SCP1 STARR-seq screen (see C and text). The upper profile is obtained from all reporter transcripts and the lower two from transcripts that either initiated in the ORI (top) or the SCP1 (bottom, see methods for details on stratification). **E**, The old and new setup of the STARR-seq plasmid (top) and STARR-seq profiles for screens using both setups in HeLa-S3 cells at a representative locus (H3K27ac data from ENCODE, Table S2). **F**, **G**, STARR-seq signal-to-noise between screens employing SCP1 or the ORI as a core-promoter over predicted enhancers for HeLa-S3 cells (F, n=39) or HCT-116 cells (G, n=27). Error bars indicate 75% confidence intervals, ** P-value < 0.01, two-sided paired t test. See Fig. S1B for an equivalent analysis over luciferase validated regions in HeLa-S3 cells. **H**, As in (E) for HCT-116 cells (H3K27ac data from Rickels et al.^57^, Table S2). **I**, Average luciferase activity for 3 cellular (AGAP1, GTSE1, IGF1R) and 2 viral (SV40, CMV) enhancers over a negative control (in log_2_ fold-change) in different reporter plasmid setups with the mCP (yellow), SCP1 (red), and ORI (green) as core-promoter. Error bars represent 1 SD across three independent transfections, n.s. not significant, * P-value < 0.05, **P-value < 0.01, ***P-value < 0.001, Fisher’s LSD test.

### STARR-seq reporter transcripts initiate in the ORI rather than in core-promoters

Similar to single-candidate luciferase assays, STARR-seq tests enhancer candidates down-stream of a core-promoter as comprehensive libraries with hundreds of millions of fragments. Candidates that function as enhancers activate transcription from the upstream core-promoter, leading to their own transcription such that enhancer activities can be measured by the candidates’ abundance among cellular RNAs^7^. To assess where reporter transcription initiates within the plasmid in STARR-seq, we mapped the initiation sites of STARR-seq reporter transcripts using a recently developed method to determine reporter mRNA 5’ ends based on deep-sequencing called STAP-seq^8^. We performed these experiments in HeLa-S3 cells, using STARR-seq libraries with two frequently used synthetic core-promoters (mCP and super core promoter 1, SCP1^9^), two endogenous core-promoters of the TTF2 and SULT1C2 genes, and one non-core-promoter control. In line with the findings for luciferase reporters^3^, the vast majority of the STARR-seq reporter transcripts (≥77.5%) initiated within the ORI and almost no initiation occurred at the different core-promoters (≤1.5%; Fig. 1B). Even SCP1, which was the only exception giving rise to 27.0% of all STARR-seq reporter transcripts, was much less esfficient compared to the ORI (59.1% of the reporter transcripts).

### ORI-derived STARR-seq reporter transcripts identify active enhancers

To test to what extent the resulting enhancer activity profiles were affected by this predominant initiation within the ORI, we processed the same samples using the standard STARR-seq protocol^7^ and mapped the reporter transcripts to the human genome (Fig. 1C). All five tested constructs show similar profiles that identify active enhancers, despite considerable background, especially at GC-rich exons. When we stratified the STARR-seq reporter transcripts for the SCP1 screen according to whether they initiated within SCP1 or the ORI, we obtained highly similar enhancer-activity profiles (Fig. 1D, see methods). This suggests that the ORI functions as an efficient and highly inducible core-promoter that responds to human enhancers.

### The ORI is an optimal core-promoter for STARR-seq and luciferase assays

To capitalize on the efficiency of the ORI as a core-promoter, we cloned STARR-seq libraries in which the ORI is used as core-promoter, placed immediately upstream of the reporter genes that contain the enhancer candidates. This should provide maximal enhancer mediated activation, avoid the presence of two potentially conflicting core-promoters on the same plasmid, and prevent the transcription and diverse splicing processes reported to occur across the ~2 kb intervening plasmid sequence^3^. Indeed, STARR-seq in HeLa-S3 cells with these constructs produced enhancer activity profiles with improved signal-to-noise for putative enhancers predicted based on chromatin features (Fig. 1E,F) and for luciferase-validated enhancers (Fig. S1B).

Similarly, using the ORI as a core-promoter improved the signal-to-noise ratio of STARR-seq screens in the unrelated colorectal cancer cell line HCT-116 (Fig. 1G,H). Importantly, these improvements were not specific to STARR-seq but applied also to single candidate luciferase assays: reporter transcription was induced up to 10-fold more strongly by cellular and up to 40-fold more strongly by viral enhancers in the ORI-based setup compared to mCP-or SCP1-based setups (Fig. 1I).

Together, these results suggest that employing the ORI as a core-promoter to assess enhancer activities improves the signal-to-noise in both single candidate luciferase assays and MPRAs such as STARR-seq. This strategy makes specific use of the ORI’s strong function as a core-promoter in mammalian cells. In addition, we also developed an alternative strategy that uses a combination of splice acceptors and polyadenylation signals to capture and suppress ORI-derived transcripts and therefore allows screens with any core-promoter of choice (Fig. S1C). Lastly, the use of ORI-less linear DNA fragments (e.g. created by PCR) or ORI-less minicircles^10–12^ and doggybones^13^ should be possible. It will be interesting to see if other low(er)-copy ORIs also have core-promoter functionality and if they allow the cloning of highly complex candidate libraries.

### DNA transfection into human cells mounts an interferon response

Reminded of the obstacles that were encountered when introducing RNA-interference from fly to mammalian cells^14^, we hypothesized that enhancer activity assays in human cells might also suffer from an innate immune response mounted against cytoplasmic DNA, which is prevalent during plasmid DNA transfection^15,16^. Most mammalian cell types and many immortalized cell lines sense cytoplasmic DNA and induce type-I-interferon (IFN-I) expression via cGAS, STING, TBK1, and IRF transcription factors^17–19^. This substantially alters gene expression, suggesting that the corresponding enhancer activities are also altered.

The expression of innate immunity genes argues, for example, that most ENCODE cell lines, which are widely used to study transcription regulation and enhancer biology^20^, have an intact cGAS/STING signaling pathway (Fig. 2). Indeed, based on the expression of interferon pathway genes, 19 ENCODE cell lines cluster into two main groups (Fig. 2A): six cell lines including HCT-116 show only low expression levels of the canonical DNA sensing pathway member IFI16 and seem to have lost cGAS or STING, suggesting that their ability to induce IFN-I expression in response to cytoplasmic DNA might be compromised. In support of this notion, five of the six cell lines were previously reported to show no or only weak upregulation of interferon stimulated genes (ISGs) upon introduction of cytosolic DNA^21–25^ (equivalent information was not available for H1-hESCs). In contrast, the majority of all cell lines (13 out of 19), including the widely-used HeLa-S3 cells, show high expression of interferon pathway genes including cGAS and STING, indicative of a functioning innate immune response to cytoplasmic DNA.

**Figure 2.**
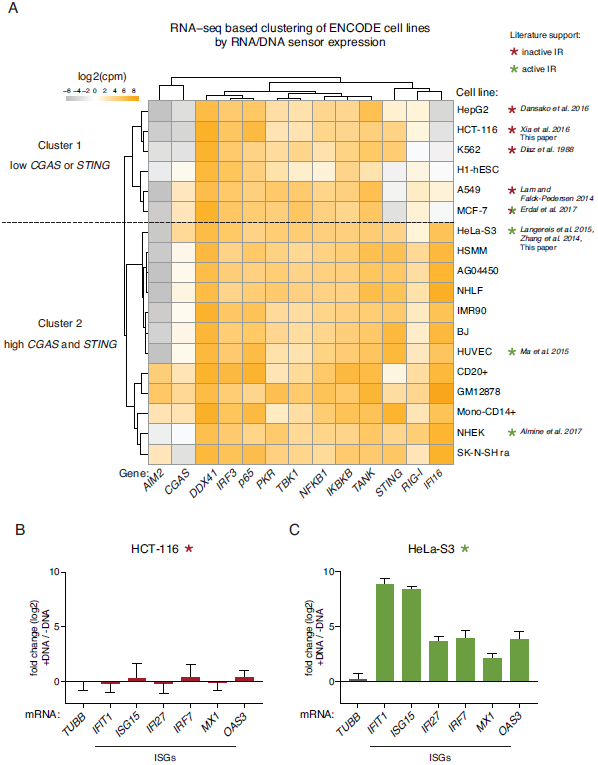
Most ENCODE cell lines are likely capable of mounting an IFN-I response to cytosolic nucleic acids **A**, Hierarchical clustering of ENCODE cell lines based on their expression profiles of genes involved in DNA-and RNA-triggered innate immunity. The column to the right indicates literature-support for (green) or against (red) a functioning cGAS/STING pathway^21–25,60–63^. **B**, **C**, qPCR-based assessment of mRNA induction after DNA transfection for canonical interferon stimulated genes (ISGs) in HCT-116 cells (B) or HeLa-S3 cells (C). Shown is the fold change in mRNA expression levels (log_2_) after DNA transfection, error bars represent 1 SD over three independent transfections.

### HeLa-S3 but not HCT-116 cells induce interferon stimulated genes

To test if HeLa-S3 cells but not HCT-116 cells induce ISGs upon DNA transfection, we determined the expression of six ISGs before and after transfecting a STARR-seq plasmid library by electroporation. In agreement with the expression of dsRNA and DNA sensors, HCT-116 cells did not induce ISGs (Fig. 2B), consistent with defective IFN-I induction. In contrast, HeLa-S3 cells strongly induced ISG expression upon DNA transfection (Fig. 2C) and ISG induction was not specific to the introduction of a STARR-seq plasmid library, but also occurred when we transfected luciferase reporter plasmids (Fig. S2A) or plasmids without any known mammalian regulatory sequences (pBluescript; Fig. S2A; see also^14^). The upregulation also did not depend on the method of plasmid delivery, as chemical transfection led to the same, up to 1000-fold, activation of ISGs (Fig. S2B).

### Enhancer activity assays in HeLa-S3 cells are dominated by IFN-I related signaling

The IFN-I-related gene induction suggests that the corresponding enhancer and promoter activities are also strongly changed. Indeed, putative enhancers proximal to canonical ISGs upregulated upon DNA transfection in HeLa-S3 cells (Fig. 2C, S2A&B) showed high activity levels in single-candidate luciferase assays, similar to strong human enhancers (Fig. 3A). Moreover, a genome-wide STARR-seq screen in HeLa-S3 cells was dominated by enhancers related to IFN-I signaling: genes proximal to the top 1000 enhancers were strongly enriched for GO terms relating to cellular immunity and IFN-I signaling (Fig. 3B, C), similar to what was recently observed in a focused STARR-seq screen testing ~21,000 promoter regions for enhancer activity^26^. This enrichment is not consistent with gene expression in unperturbed HeLa cells, because genes in these categories are not particularly highly expressed^27^.

**Figure 3.**
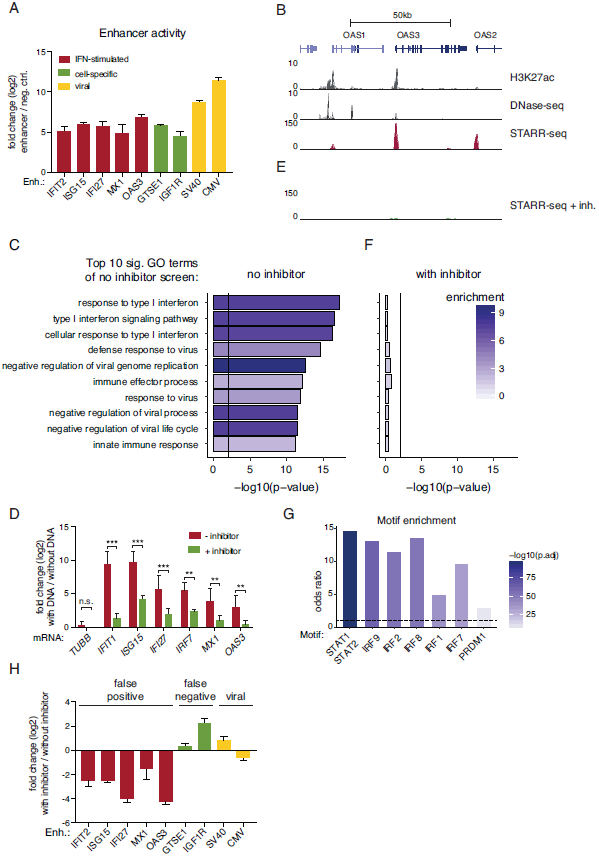
TBK1/IKK*ε* and PKR inhibition prevents dominant false-positive IFN-I related enhancer signals **A**, Average enhancer activity measured as luciferase mRNA induction over negative control (log_2_) as measured by qPCR in reporter assays employing the indicated enhancers. Error bars represent 1 SD across three independent transfections. **B**, Representative STARR-seq enhancer activity profiles over a canonical ISG locus (GRCh37 Refseq genes indicated above) for genome-wide HeLa-S3 screens without inhibitors against TBK1/IKK/PKR (H3K27ac and DHS data from ENCODE, Table S2). **C**, The 10 most significantly enriched GO terms for genes proximal to the top 1000 peaks in a HeLa-S3 STARR-seq screen. Shown are P-values (Fisher’s exact test) and fold-enrichments (shades of purple). The same terms were assessed for the TSSs proximal to the top 1000 peaks from the TBK1/IKK/PKR-inhibitor-treated screen (F). **D**, qPCR-based assessment of ISG-mRNA induction after DNA transfection in TBK1/IKK/PKR-inhibitor-treated vs. non-treated HeLa-S3 cells. Error bars represent 1 SD across three independent transfections, n.s. not significant, **P-value < 0.01, ***P-value < 0.001, Fisher’s LSD test. **E**, Same as B for a genome-wide HeLa-S3 screen with inhibitors against TBK1/IKK/PKR. **F**, The same terms that were assessed in (C) for the TSSs proximal to the top 1000 peaks from a TBK1/IKK/PKR-inhibitor-treated screen. **G**, Odds ratios (FDR-adjusted P-values < 10-5, Fisher’s exact test) of indicated transcription factor motifs in STARR-seq enhancers 5-fold downregulated upon TBK1/IKK/PKR-treatment (FDR-adjusted P-value < 0.001) vs. unchanged enhancers (within +/-1.5-fold change upon treatment). **H**, Average luciferase activity fold change of luciferase mRNA expression in reporter assays employing the indicated enhancers in cells treated without PKR/TBK1 inhibitors over with inhibitors (log_2_). Error bars represent 1 SD across three independent transfections.

### TBK1/IKK *ε* and PKR inhibition prevents dominant false-positive IFN-I related enhancer signals

Cytoplasmic DNA leads to IFN-I induction via the key signaling kinases TANK binding kinase 1 (TBK1) and I*к*B kinase *ε* (IKK*ε*), which activate IRF3^28^. Similarly, double stranded RNA (dsRNA) that can arise at transfected plasmids is sensed by dsRNA-activated protein kinase (PKR) and can affect transgene expression^29–31^. The inhibition of these key signaling kinases should therefore ameliorate the changes of gene expression and enhancer activities described above. Indeed, treating HeLa-S3 cells during plasmid transfection with the TBK1/IKK*ε* inhibitor BX-795^32,33^ and the PKR inhibitor C16^34^ prevents the strong induction of ISGs observed after plasmid transfection in HeLa-S3 cells (Fig. 3D).

Moreover, enhancers proximal to canonical ISGs that were previously among the strongest STARR-seq signals genome-wide, were only detected at background levels in an inhibitor-treated STARR-seq screen (Fig. 3E, compare to Fig. 3B). Furthermore, genes near the top 1000 peaks no longer showed the enrichment in interferon-signaling-related GO categories observed without inhibitor treatment (Fig. 3F, compare to Fig. 3C), indicating that TBK1/IKK/PKR inhibition removes these dominant false-positive signals. Consistently, STARR-seq peaks that lost activity upon TBK1/IKK/PKR inhibition (≥ 5-fold down-regulation, P-value < 0.001) were next to IFN-I signaling-related genes (Fig. S3A) and were highly enriched in binding motifs of IRF and signal transducer and activator of transcription (STAT) transcription factors, known to be involved in ISG induction (Fig. 3G). Finally, luciferase assays confirmed that enhancers proximal to ISGs showed strongly reduced activity in cells treated with TBK1/IKK/PKR inhibitors during plasmid transfection (Fig. 3H). Overall, this demonstrates that plasmid-induced IFN-I-related false positive enhancer activities can be prevented by TBK1/IKK/PKR inhibition.

### A genome-wide set of IFN-I related enhancers

The comparison of the genome-wide STARR-seq screens with and without TBK1/ IKK/PKR inhibitors also defines a genome-wide set of IFN-I related enhancers and their respective induction strengths, some of which were more than 100 fold (Fig. S3D, Table S3). Interestingly, many of these predominantly promoter-proximal enhancers were pre-marked by H3K27ac even when uninduced in unperturbed HeLa-S3 cells (Fig. S3D), which might be a general feature of rapidly inducible enhancers downstream of stress signaling pathways^35^. This genome-wide set of IFN-I related enhancers (Table S3) should be a valuable resource for the study of IFN-I mediated transcriptional regulation.

### TBK1/IKK/PKR inhibition prevents false-negative signals and improves signal-to-noise

Interestingly, of the enhancers we tested individually in luciferase assays, the activity of two endogenous HeLa-S3 cell enhancers and one viral enhancer was increased after TBK1/IKK/PKR inhibition (Fig. 3H). This suggests that IFN-I-mediated transcriptional regulation might also repress bona fide enhancers, leading to an underestimation of their activities or false negative results if TBK1/IKK/PKR inhibitors are not used. Indeed, the STARR-seq signal-to-noise ratio in inhibitor-treated HeLa-S3 cells increased substantially in comparison to untreated cells (Fig. S3B). In contrast, inhibition of PKR and TBK1/IKK did not change the STARR-seq signal-to-noise ratio in HCT-116 cells, which do not induce ISGs in response to DNA transfection (Fig. S3C). This indicates that the improvement in signal-to-noise is specific to effects related to the interferon response and that inhibitor treatment does not otherwise impact enhancer activities.

### STARR-seq enhancers are mostly intergenic or intronic and are enriched in enhancer-associated chromatin states

Overall, the genome-wide STARR-seq screen using the new screening setup and TBK1/IKK/PKR-inhibitors yielded 9,613 peaks, of which 2,508 have a corrected enrichment of ≥10-fold and 209 of ≥50-fold. The enhancer activity profiles are highly similar between independent replicates (PCC=0.98).

Almost half of all peaks are in intergenic regions (48.3%) and 43.2% are within introns (Fig. S4A). The peaks highly significantly overlap with regions that exhibit enhancer-or promoter-associated chromatin states according to chromHMM^36^: all 9,613 STARR-seq peaks are enriched 20.5-fold in the strong enhancer state *Enh*, 6.7-fold in the weak enhancer state *EnhW* and 6.6-fold in the active promoter state *TSS*, compared to these states’ genomic abundance (Fig. 4A). The enrichment for the *Enh* state was as high as 40.3-fold for the top 500 peaks and still ~ 9-fold for the weakest called peaks, suggesting that even peaks below the threshold used for peak calling might be functional *in vivo*, albeit with weak effects on transcription activation (Fig. S4B, Table S3). Furthermore, peaks that are accessible in HeLa-S3 cells according to ENCODE DNase-seq (42.3%, Table S2) align precisely with characteristic enhancer features (Fig. 4B).

**Figure 4.**
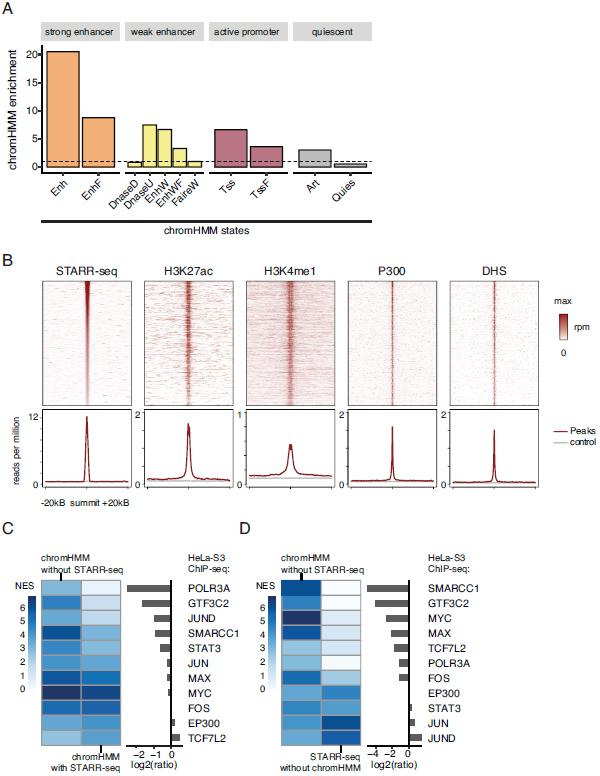
STARR-seq enhancers are enriched in enhancer-associated states**A**, Enrichment of enhancer relevant ChromHMM states^36^ within STARR-seq enhancers (dotted line indicates no enrichment (1)). **B**, Heatmaps of reads per million (top) and average signal (bottom) of STARR-seq, H3K27ac, H3K4me1, P300 and DHS signal for STARR-seq enhancers accessible in HeLa-S3 cells. The average signal is compared to a set of random control regions (grey, see methods). **C**, Normalized enrichment scores for different HeLa-S3 ChIP-seq datasets (NES, i-cisTarget^37^) for chromHMM strong enhancers (’Enh’) with our without STARR-seq support (as indicated) and ratios between NES (right panel, log_2_). **D**, Normalized enrichment scores (NES, i-cisTarget^37^) for different HeLa-S3 ChIP-seq datasets for chromHMM strong enhancers (’Enh’) without STARR-seq support and open STARR-seq enhancers that do not overlap chromHMM strong enhancers (’Enh’) and ratios between NES (right panel, log_2_).

### STARR-seq-negative enhancer-candidates are associated with Pol III transcription

ChromHMM *Enh* regions that do not show any activity in STARR-seq and might not function as enhancers or do so only weakly are enriched for ENCODE-defined binding sites of RNA polymerase III (Pol III) and its general transcription factor 3C (Fig. 4C, enrichments assessed with i-cisTarget^37^). Pol III typically transcribes non-coding genes such as tRNAs from promoters that are independent of enhancer-like upstream regulatory regions^38^. However, regions with Pol III occupancy can bear chromatin marks reminiscent of Pol II enhancers^39^, which presumably explains their annotation by ChromHMM. Of all chromatin-related datasets considered by i-cisTarget that are enriched in ChromHMM regions with or without STARR-seq signals, Pol III and GTF3C2 were the most differentially enriched. Others, including the transcription factors Jun, Max, Myc, or Fos were enriched to similar extents in ChromHMM regions with or without STARR-seq signals (EP300 and TCF7L2 were slightly more enriched in ChromHMM regions with STARR-seq support). Open STARR-seq enhancers that do not overlap chromHMM *Enh* regions show similar enrichments for different TFs and an even slightly higher enrichment for the TFs JUN and JUND as well as the transcriptional activator EP300, as expected for bona fide enhancers (Fig. 4D).

### A substantial fraction of enhancers is silenced at the chromatin level

One of the advantages of ectopic assays such as STARR-seq is their ability to assess the enhancer activities of DNA sequences that are able to strongly activate transcription yet are silenced endogenously at the chromatin level^7^. While as expected a large fraction of the STARR-seq enhancers were accessible in their endogenous genomic locations according to ENCODE DNase-seq (56.2% of the top 500 and 42.3% of all 9,613 enhancers, Table S2), 57.7% of STARR-seq enhancers were not accessible, i.e. closed, and thus likely actively silenced (Fig. 5A). This included DNA sequences that can function as very strong enhancers in HeLa-S3 cells, such as peak 89 in the *CWC27* locus or peak 384 in the *HMX1* locus (Fig. 5B).

**Figure 5.**
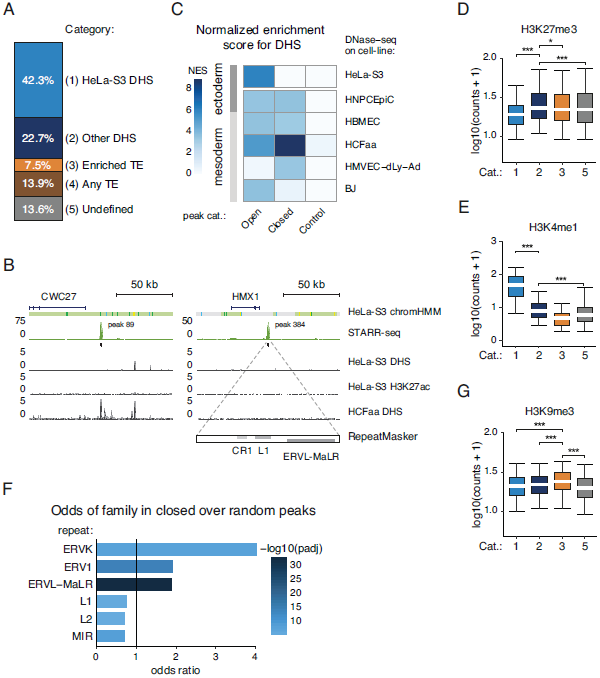
STARR-seq identifies enhancers that are silenced endogenously **A**, Percentages of STARR-seq enhancers that have significant DNase-seq signal in HeLa-S3 cells (P-value < 0.05, binomial test), are accessible in other enriched cell types, contain repetitive elements from three enriched repeat families (see Fig. 5G), contain other repetitive elements, or none of the above (undefined). **B**, Enhancer activity profiles over two gene loci (indicated above; DHS and H3K27ac data from ENCODE, Table S2), representative of category 2 (CWC27, left panel) and 3 (HMX1, right panel). The right panel includes the RepeatMasker track, displaying elements of the indicated repeat families within the STARR-seq peak above. **C**, Normalized enrichment scores (NES, i-cisTarget^37^) for ENCODE DNase-seq datasets within STARR-seq enhancers that are open or closed in HeLa-S3 cells (P-value < 0.05, binomial test). NES scores for random regions are shown as control. **D**, **E**, **G**, Boxplots of H3K27me3 (D), H3K4me1 (E) and H3K9me3 (G) read coverage in log_10_ (counts + 1) for STARR-seq enhancers of the categories defined in (B). Lower whisker: 5^th^ percentile, lower hinge: 25^th^ percentile, median, upper hinge: 75^th^ percentile, upper whisker: 95^th^ percentile. **F**, Odds ratios (FDR-adjusted P-value < 0.05, Fisher’s exact test) of indicated transposable elements in STARR-seq enhancers inaccessible in HeLa-S3 cells (DHS, P-value < 0.05, binomial test) vs. 1×10^6^random control regions.

### Closed STARR-seq enhancers are open in other cell types and H3K27me3 marked

Interestingly, the closed enhancers are strongly enriched in DNase I hypersensitive regions of several ENCODE cell lines according to i-cisTarget^37^. These enrichments are partly stronger than the enrichments for open HeLa-S3 enhancers (e.g. for HMVEC or HCFaa cells; Fig. 5C). Overall, DNase I hypersensitivity in the five non-HeLa cell types accounts for 39.3% of the closed HeLa-S3 enhancers (Fig. 5A, see Table S2).

Moreover, in HeLa-S3 cells, these closed enhancers were enriched for the H3K27me3 mark compared to accessible peaks or peaks containing repetitive elements (Fig. 5D, see below), consistent with Polycomb-mediated repression^40^. Interestingly, they were marked by H3K4me1 (albeit not to the same extent as accessible enhancers; Fig. 5E) suggesting that they might be recognized as enhancers as previously observed in flies^7^. This is also consistent with reports from both flies and mammals that the H3K4me1 mark labels enhancers independently of their activity^41–43.^

### Closed STARR-seq peaks are enriched for TEs and are H3K9me3-marked

Many (39.8%) of the remaining closed peaks (24.1% of all closed peaks) contained repetitive elements annotated by RepeatMasker^44^ (Fig. 5A). Interestingly, while LINE elements (L1 and L2) were depleted overall within closed peaks, 3 families of endogenous retroviruses were highly enriched and elements from just these 3 families overlapped with 13.0% of all closed peaks (Fig. 5F). The non-accessible peaks that overlapped elements of the enriched repeat families exhibited a significantly higher signal of H3K9me3 than other regions (Fig. 5G). This finding and the observation that such elements can be co-opted for transcriptional regulation ^45,46^ and are induced when DNA methyltransferases or histone de-acetylases are inhibited^47^, suggest that these sequences indeed constitute active regulatory regions that are repressed by heterochromatin proteins and H3K9 methyltransferases^48^.

## Discussion

The past years have seen tremendous progress in the prediction of transcriptional enhancers in mammalian genomes based on chromatin properties and non-coding transcription that correlate with enhancer activities^2,49^. The direct functional assessment of distal enhancer activities by reporter assays has therefore become increasingly important and is a major aim of large consortia efforts^50^. Functional tests with reporter assays are also important for many mechanistic studies, particularly because such reductionist approaches can exclude confounding effects present in complex endogenous loci and constitute direct tests of a defined DNA sequence’s sufficiency for distal transcriptional activation and its strength.

Here, we show that two previously reported effects – the core-promoter function of a bacterial ORI and the IFN-I response triggered by cytoplasmic DNA – confounds enhancer activity assays in mammalian cells. In fact, our results indicate that previous approaches suffering from these problems might have substantially underestimated enhancer activities and could have missed up to 75% of all enhancers (Fig. S4C). The problem related to IFN-I induction is reminiscent of the early days of RNAi in mammalian cells, which – in contrast to Drosophila cells – mount an interferon response in the presence of the long double-stranded RNA used for RNAi^14^. We provide simple means to prevent the interferon response triggered by cytoplasmic DNA or dsRNA and – importantly – to assess if a particular cell line is likely sensitive to these stimuli. In fact, we expect most if not all primary cells and the majority of ENCODE cell lines to mount an interferon response to cytoplasmic DNA and/or dsRNA (Fig. 2A) such that the proposed tests and counter-measures will be crucial for all future enhancer activity studies. As the INF-I induced enhancers are predominantly promoter-proximal and might also function as promoters^26^, promoter-activity assays (e.g. ref. 51) or MPRAs with candidates positioned upstream of the reporter gene or barcode^2^ should also be affected. We anticipate that similar considerations apply to other signaling pathways in mammalian cells that might be triggered by plasmid transfection. Depending on the cell type, DNA transfection might for example trigger a DNA damage response via p53 (particularly in primary cells – p53 is often inactivated in cancer cells^52^) or an inflammatory response via NFkB^53^.

The methods presented here overcome two key problems associated with plasmid transfection during plasmid-based enhancer-activity assays. For cells that can be transfected, plasmid-based assays are highly efficient and comparatively cheap - in contrast, for example, to assays based on virus-mediated infection that typically require substantially higher effort and increased biosafety levels. Furthermore, integrating viruses often show systematic preferences to insert into open chromatin near transcriptionally active genes^54–57^, such that enhancer activity measurements might be influenced by endogenously active enhancers and promoters near the integration sites. While a recent paper found no major differences between integrated and non-integrated assays (ref 58, Fig. 3 therein) and the influence of individual integration sites might be averaged out, systematic preferences should cause systematic biases that are difficult to foresee and control.

The genome-wide STARR-seq screens in one of the most widely used model cell lines, HeLa-S3 cells, highlight the power and importance of systematic genome-wide enhancer identification using screens based directly on enhancer activity in human cells: it reveals thousands of human enhancer sequences with activities approaching the strengths of strong viral enhancers several hundred-fold over background. Similar to previous results in flies, many of even the strongest enhancers are silenced in their endogenous loci at the chromatin level. These enhancers are inaccessible to predictions based on chromatin features, yet constitute attractive examples to study mechanisms of chromatin-mediated repression. The genome-wide enhancer activity profiles in the presence or absence of IFN-I signaling identify strongly IFN-I responsive enhancers and are a valuable resource for future studies of IFN-I signaling and transcriptional regulation more generally.

The tools and protocols presented here should be applicable to all episomal reporter assays in mammalian cells used to assess enhancer activities of individual enhancers or on a genome-wide level. They are of central importance to all ongoing efforts that validate enhancer predictions or, for example, study the functional impact of single-nucleotide polymorphisms with such assays, particularly in cells with intact innate immune signaling. Given the exemplary results in HeLa-S3 and HCT-116 cells, we anticipate that both, the testing of individual candidates and genome-wide screens with the tools and protocols presented here will become a central component of our efforts to identify all gene regulatory elements of the human genome and understand how their sequences encode cell type-specific gene expression.

## Acknowledgements

We thank Thomas Decker and Gijs Versteeg (Max F. Perutz Laboratories & University of Vienna), Stein Aerts (VIB-KU Leuven), Petr Svoboda (Institute of Molecular Genetics of the ASCR) and Peter Andersen (IMBA) for helpful discussions. Deep sequencing was performed at the VBCF Next-Generation Sequencing Unit (*http://vbcf.ac.at*). F.M. was supported by an EMBO long-term fellowship (EMBO ALTF 491–2014). Research in the Stark group is supported by the European Research Council (ERC) under the European Union’s Horizon 2020 research and innovation programme (grant agreement no. 647320) and by the Austrian Science Fund (FWF, F4303-B09). Basic research at the IMP is supported by Boehringer Ingelheim GmbH and the Austrian Research Promotion Agency (FFG).

## Author Contributions

F.M. and L.M.B. are shared first authors, A.R.W and C.N. are shared second authors. F.M., L.M.B., C.N., M.R., M.P., R.R.C., K.S. and C.D.A. performed experiments. L.M.B. designed and cloned all STARR-seq and luciferase vectors. A.R.W. and F.M. performed the computational analysis with the help of M.A.Z.; V.H. and T.K. analyzed the ORI sequence. F.M. and A.S. wrote the manuscript with help from all authors. A.S. supervised the project.

## Supplemental figures

**Figure S1.**
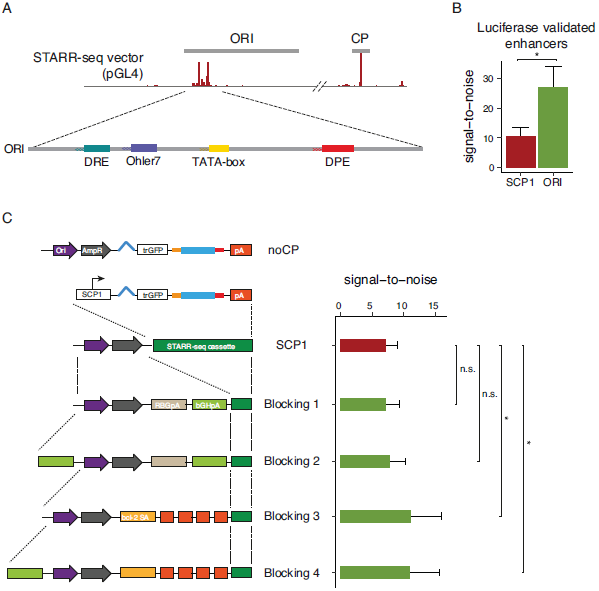
ORI initiation and blocking of ORI derived transcripts **A**, The core-promoter motif content is shown for the ORI of the pGL4 family. The track above is taken from Fig. 1B, panel for SCP1. **B**, STARR-seq signal-to-noise between screens employing SCP1 or the ORI as a core-promoter over 10 luciferase validated enhancers (4 positive, 6 negative, see methods for details). Error bars indicate 75% confidence intervals, *P-value < 0.05, two-sided paired t test. **C**, Alternative strategy to block ORI-derived transcripts based on the insertion of poly-adenylation sites and splice acceptors downstream of the ORI. Note that the introduction of poly-adenylation sites alone – as present for example in pGL3 and pGL4 – are ineffective due to extensive splicing of the ORI-derived transcript (compare blocking constructs 1 and 2 to 3 and 4)^3^. The right panel depicts signal-to-noise for the indicated constructs over predicted enhancers (see methods). Error bars indicate 75% confidence intervals, *P-value < 0.05, two-sided paired t-test. Alternatively, one might create plasmids that do not contain the ORI and thus only a single core-promoter that one should be able to choose freely. A family of vectors designed to remove prokaryotic parts of the plasmid after amplification in bacteria (minicircles^10–12^) or in vitro synthesis (doggybones^13^) could be such alternatives, as might be different lower copy ORIs. Alternatively, one could screen linear DNA fragments that lack the ORI (e.g. PCR amplicons of candidate fragments). However, it remains to be seen if any of these alternatives are compatible with the construction of highly complex libraries.

**Figure S2.**
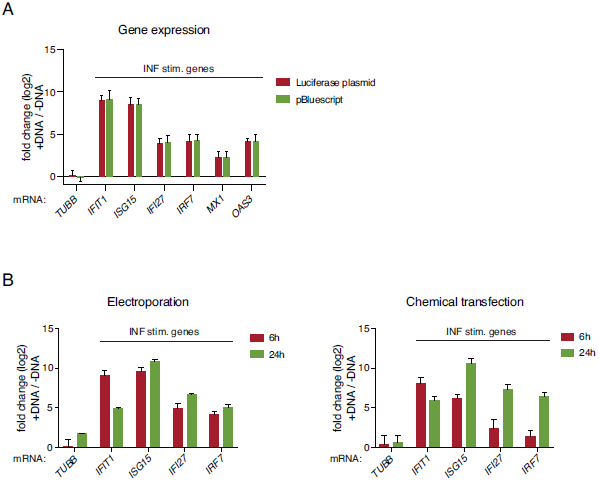
IFN-I induction is independent of plasmid type or transfection method **A**, **B**, qPCR-based assessment of mRNA induction of ISGs in HeLa-S3 cells electroporated with different DNA sources (A) and by different means of transfection (B). In B, RNA extraction was performed at two different time points to account for the different kinetics of DNA delivery between electroporation and chemical transfection. In all cases, fold change in mRNA expression levels (log_2_) after transfection is shown; error bars represent 1 SD over three independent transfections.

**Figure S3.**
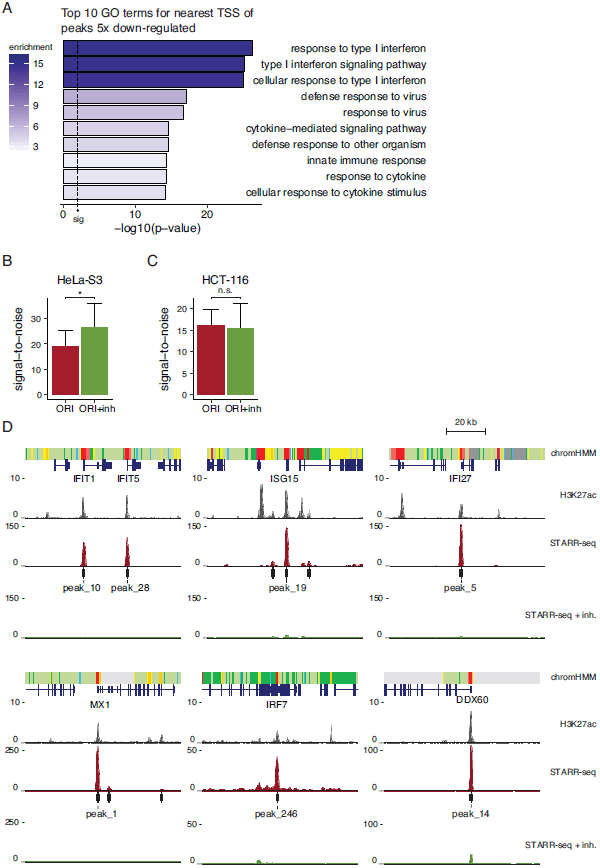
TBK1/IKK/PKR inhibition prevents false-positive and false-negative errors **A**, Top 10 GO terms (ranked by P-value, Fisher’s exact test) and their enrichments among genes proximal to enhancers that are at least 5-fold down-regulated upon PKR/TBK1 inhibitor treatment (FDR-adjusted P-value < 0.001). **B**,**C**, Average signal-to-noise of STARR-seq screens in cells with (green) or without (red) TBK1/IKK/PKR-inhibitor treatment over predicted enhancers for HeLa-S3 cells (B, n = 39, see methods) or HCT-116 cells (C, n = 27). Error bars indicate 75% confidence intervals, n.s. not significant, *P-value < 0.05, two-sided paired t test. **D**, Representative STARR-seq enhancer activity profiles for canonical ISGs. Note that their differential activity in screens without (red) and with (green) TBK1/IKK/PKR inhibition is a good indicator for their inducibility by IFN-I. The peak ranks are listed below each strong enhancer indicating that these regions are among the most highly active in STARR-seq screens performed without inhibitors.

**Figure S4.**
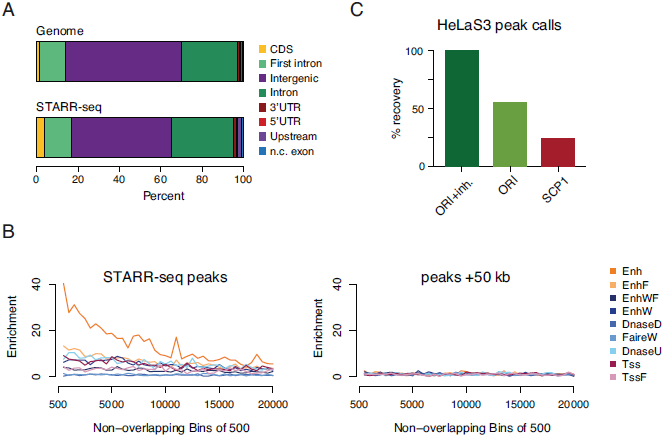
STARR-seq enhancers are mostly intergenic or intronic and are enriched in enhancer-associated chromatin states. **A**, Average percentages of genomic annotations for the human genome (top) and STARR-seq peaks (bottom). CDS = coding sequence, 3’ / 5’ UTR = 3’ / 5’ untranslated region, n.c. exon = non-coding exon, upstream = −2 kb from the TSS. **B**, Enrichment of different ChromHMM states^36^within non-overlapping bins of 500 STARR-seq enhancers (ranked by corrected fold-enrichment, left) and shifted control regions (+50kb, right). **C**, Percent recovery of peak calls in focused STARR-seq screens in HeLa-S3 cells employing the ORI or SCP1 as core-promoters as a fraction of peak calls in the ORI setup with inhibitor treatment.

## Methods

The methods section is accompanied by detailed laboratory protocols for STARR-seq library cloning and screening. We further provide detailed protocols for qPCR based testing of induction of an interferon response and qPCR based reporter assays of luciferase transcripts. All files are available at http://starklab.org/data/muerdter_boryn_2017/. All plasmids are available from Addgene.

## Experimental/ Laboratory Methods

### Cell culture

Human HeLa-S3 were purchased from American Type Culture Collection (ATCC; cat. no. CCL-2.2). HCT-116 cells were a kind gift from the Zuber lab (IMP), originally purchased from ATCC (cat. no. CCL-247). Cells were cultured in DMEM (Gibco; cat. no. 52100-047), supplemented with 10% heat-inactivated FBS (Sigma; cat. no. F7524) and 2 mM L-glutamine (Sigma; cat. no. G7513) at 37°C in a carbon dioxide (CO_2_) enriched atmosphere (95% air and 5% CO_2_). Cells were harvested at 80% confluency by removing the growth medium, washing with 1x PBS, treating with 0.25% Trypsin-EDTA (Gibco; cat. no. 25200-056) until dispersion of the cell layer and resuspension in complete medium.

### Transfection by electroporation

Cells were electroporated using the MaxCyte STX Scalable Transfection System, as recommended by the manufacturer. Briefly, harvested cells were pelleted, washed with 5 ml MaxCyte HyClone electroporation buffer (MaxCyte; cat. no. EPB5), resuspended in HyClone electroporation buffer and mixed with plasmid DNA. Cells were electroporated at a density of 1×10^7^ cells per 100 μ;l and 20 μg of DNA in HyClone electroporation buffer in an OC-100 processing assembly (MaxCyte; cat. no. GOC1) or 4×10^7^ cells and 80 μg DNA in 400 μl HyClone buffer in an OC-400 processing assembly (MaxCyte; cat. no. GOC4), applying the pre-set cell type-specific electroporation programs provided by the manufacturer (i.e. Hela or HCT 116 protocols). After electroporation, cells were immediately transferred to a pre-warmed tissue culture flask allowing them to recover as cell drop without addition of medium for 30 minutes at 37°C. After this essential recovery phase, cells were resuspended in the appropriate volume of complete growth medium to be plated at 70% confluency.

### Chemical transfection

8×10^5^ HeLa-S3 cells were plated in 8 ml complete growth medium in T-25 flasks (Invitrogen; cat. no. 156367) 24 h prior to chemical transfection. To prepare the transfection mix, 8.8 μg of STARR-seq library was diluted in a total volume of 409 μl medium followed by the addition of 31 μl FuGENE HD transfection reagent (3.5:1 reagent to DNA ratio, Promega; cat. no. E2312). After 5 minutes incubation to allow complex formation, 400 μl complexes were dropwise added onto the cells. After 6h or 24h, total RNA from all transfected cells was extracted using the Qiagen RNeasy mini prep kit (Qiagen; cat. no. 74104).

### Inhibitor treatment

The PKR (C16, Sigma; cat. no. I9785-5MG) and TBK1/IKK inhibitors (BX-795 hydrochloride, Sigma; cat. no. SML0694-5MG) were added to the electroporated cells directly after resuspension in complete growth medium at a final concentration of 1 μM per inhibitor. Note: Both inhibitors are kept at −20°C as 10 mM stocks in DMSO.

### Mapping of STARR-seq transcript initiation sites by STAP-seq

STAP-seq was performed as described before^8^. In brief, 50 μg of DNaseI-treated mRNA from cells electroporated with a STARR-seq library was treated with 25 μl of Calf Intestinal Alkaline Phosphatase (CIP) (NEB; cat. no. M0290L) for 1.5h at 37°C. The CIP-treated RNA was then purified using the Qiagen RNeasy MinElute reaction clean-up kit (cat. no. 74204), according to the manufacturer’s protocol, with beta-Mercaptoethanol (Sigma-Aldrich; cat. no. 63689) supplemented RLT buffer. The CIP-treated RNA was then treated with 0.05 l Tobacco Alkaline Phosphatase (TAP, Epicentre; discontinued, now available as Cap-Clip Acid Pyrophosphatase (cat. no. C-CC15011H) from CELLSCRIPT) per 1 g RNA to remove the 5 cap of all 5 -capped RNA species and purified with Agencourt RNAClean XP beads (Beckman Coulter; cat. no A66514) at a beads-to-RNA ratio of 1.8. We ligated 10 μM RNA adapter (STAP_adapter, see Table S1) to the 5’ ends of each 1 μg TAP-treated mRNA using 0.2 μl T4 RNA Ligase 1 (NEB; cat. no. M0204L) for 16h at 16°C. The RNA was purified with Agencourt RNAClean XP beads at a beads-to-RNA ratio of 1.0.

First strand cDNA synthesis was performed on the total amount of adapter ligated RNA. Per reaction 2.5–5 μg adapter ligated RNA was reverse transcribed with 1 l of Invitrogen’s Superscript III (50 °C for 60 min, 70 °C for 15 min; cat. no. 18080085) and a reporter-RNA-specific primer (STAP_GSP, Table S1). Five reactions were pooled and 1 l of 10 mg/ml RNaseA was added (37 °C for 1 h) followed by Agencourt AMPureXP DNA bead purification (ratio beads/RT reaction 1.8). We amplified the total amount of reporter cDNA for Illumina sequencing. For focused libraries, we performed two PCR reactions using the KAPA real-time library amplification kit (KAPA Biosystems, cat. no. KK2702) according to the manufacturer’s protocol, with STAP_fwd (Table S1) as forward primer and one of NEBNext Multiplex Oligos for Illumina (NEB; cat. no. E7335 or E7500) as reverse primer. PCR products were purified with AMPureXP DNA beads (ratio beads/PCR 1.25).

### STARR-seq

*Related to supplemental protocols: “human STARR-seq Library Preparation Protocol” and “human STARR-seq Protocol”*

### Human STARR-seq Screening Vectors

All STARR-seq screening vectors are based on the original human STARR-seq vector^7^ with the following changes: The GFP coding sequence is truncated, the synthetic intron^64^ is replaced with a chimeric intron^65^, and the core-promoter is replaced with a panel of different minimal promoters (see main text and supplemental Table S1).

To allow for the use of a specific core-promoter in the presence of the origin-of-replication (ORI), we generated four screening vectors containing the SCP1 core-promoter and the following changes (see Fig. S1): Blocking variant 1 contains the polyadenylation signal sequence of the rabbit B-globin gene RBGpA^66^ and the bovine growth hormone polyadenylation signal bGHpA^67^ in the SpeI restriction site. Blocking Variant 2 additionally contains the bGHpA sequence upstream of the ORI in the PciI restriction site. To prevent splicing over the polyadenylation signals, we constructed two additional variants: blocking variant 3 contains the human bcl-2 splice acceptor and four late SV40 polyadenylation sites at the SpeI cutting site^67^. Blocking variant 4 additionally contains a bGHpA sequence at the PciI restriction site.

To use the ORI as a core-promoter we cloned the reverse complementary sequence of the ampicillin resistance cassette followed by the ORI sequence upstream of the chimeric intron using the PciI/PspOMI restriction sites. All sequences and cloning sites for these changes can be found in Table S1.

### Cloning of STARR-seq plasmid libraries

Focused STARR-seq libraries were generated from Bacterial Artificial Chromosome (BAC) DNA (see Table S1, BAC mixes). BAC insert 1 was cloned into STARR-seq screening vectors that harbor different core promoters. BAC insert 2 was used to clone the libraries for all other screens. To generate the library inserts, BAC DNA was extracted using the Qiagen Large-Construct kit (cat. no. 12462), pooled and sheared by sonication (Covaris S220, duty cycle 2%, intensity 4, 200 cycles per burst, 15 seconds; target size: 1000bp-1500bp) followed by the library insert generation protocol, as described previously (Arnold et al. 2013). In brief, to 5 μg of size-selected BAC DNA, Illumina Multiplexing Adapters (Illumina; cat. no. PE-400-1001) were ligated using the NEBNext DNA Library Prep Reagent Set for Illumina (NEB; cat. no. E6000L) following manufacturer’s instructions, except for the final PCR amplification step. 10 PCR reactions (98°C for 45s; followed by 10 cycles of 98°C for 15s, 65°C for 30s, 72°C for 45s) with 1 μl adapter-ligated DNA as template were performed, using the KAPA Hifi Hot Start Ready Mix (KAPA Biosystems; cat. no. KK2602) and primers (in-fusion_fwd & in-fusion_rev, see Table S1) which add a specific 15nt extension to both adapters for directional cloning using recombination (Clontech In-Fusion HD; cat. no. 639650). The 10 PCR reactions were pooled, purified, and size selected on a 1% agarose gel. The purified PCR products were recombined to the STARR-seq screening vector in a total of four In-Fusion HD reactions. After pooling they were ethanol precipitated and resuspended in 12.5 μl EB [10mM Tris-HCl, ph 8]. 5 aliquots (20 μl each) of MegaX DH10B Electrocompetent Cells (Invitrogen; cat. no. C640003) were transformed with 2.5 μl DNA each, according to the manufacturer’s protocol. After 1h recovery, the transformed bacteria were transferred to 4 l selection medium (LB medium; 100 μg/ml Ampicillin) and grown in liquid culture. Bacterial cultures were harvested at an OD_600_ of 2.0-2.5. The STARR-seq plasmid libraries were extracted using the Plasmid Plus Giga Kit (Qiagen; cat. no. 12981).

To generate genome-wide STARR-seq libraries we followed the same protocol with the following changes: We used 15 μg of size-selected genomic DNA (1000bp-1500bp) for library insert generation, 30 PCR reactions for library insert amplification, 20 In-Fusion HD reactions for library cloning and transformation of 25 aliquots of Invitrogen MegaX DH10B Electrocompetent Cells. The bacteria culture was grown in 24 liters of LB.

### STARR-seq library electroporation

For focused BAC screens, we electroporated 8#x00D7;10^7^ cells per screen, for genome-wide STARR-seq libraries we electroporated 8°108 cells per screen. For details regarding transfection by electroporation using the MaxCyte STX transfection system, refer to section ‘Transfection by electroporation’. Genome-wide STARR-seq screens in HeLa-S3 cells were done in duplicate (inhibitor screens) or quadruplicate (non-inhibitor screens). Reads from independent replicates were first assessed for reproducibility and then pooled for all further analysis.

### RNA isolation

6h after electroporation, total RNA was extracted using the RNeasy Maxi kit (Qiagen; cat. no. 75162), followed by polyA+ RNA isolation using Invitrogen Dynabeads Oligo(dT)_25_ (scaling up the manufacturer’s protocol accordingly; cat. no. 61005) and DNase treatment with Ambion Turbo DNase (cat. no. AM2239) at a concentration of at most 200 ng/μl for 30 minutes (min) at 37°C. The reactions were then subjected to Qiagen RNeasy MinElute reaction clean-up (cat. no. 74204), for Turbo DNase inactivation and RNA concentration.

### Reverse transcription

First strand cDNA synthesis was performed with Invitrogen Superscript III (50°C for 60 min, 70°C for 15min; cat. no. 18080085) using at most 5 μg of polyA+ RNA per reaction and a reporter-RNA specific primer (STARR_GSP, see Table S1) in a total of 10-20 reactions for focused (BAC) STARR-seq screens and 40-60 reactions for genome-wide STARR-seq screens. Five reactions were pooled and 1 μl of 10 mg/ml RNaseA was added (37°C for 1h) followed by cDNA purification using Agencourt AMPureXP DNA beads at a beads-to-cDNA ratio of 1.8.

### Amplification of cDNA

The cDNA was PCR amplified as described before (Arnold et al. 2013) using the KAPA Hifi Hot Start Ready Mix with the following changes. The first PCR step (98°C for 45s; followed by 15 cycles of 98°C for 15s, 65°C for 30s, 72°C for 70s) of the 2-step nested PCR strategy was performed using human STARR-seq specific primers (junction_fwd, junction_rev, see Table S1), as the forward primer needs to span the splice junction of the chimeric intron. To guarantee maximal complexity, the entire cDNA was subjected to PCR amplification. For each reverse transcription reaction we performed one PCR reaction. PCR products were purified by Agencourt AMPureXP DNA beads at a beads-to-PCR ratio of 0.8. The purified PCR products from the first PCR step served as a template for the second PCR step (98°C for 45s; followed by 6-15 cycles of 98°C for 15s, 65°C for 30s, 72°C for 45s; primers PE1.0 as forward primer and MP2.0 or TruSeq idx primers as reverse primer, see Table S1) with the KAPA Hifi Hot Start Ready Mix and NEBNext Multiplex Oligos for Illumina (cat. no. E7335L). For focused STARR-seq screens, we set up 2 reactions, for genome-wide STARR-seq screens we set up 10 reactions. PCR products were purified using SPRIselect beads (Becker Coulter, cat. no. B23318) at a beads-to-PCR ratio of 0.5. The purified PCR products were pooled and the DNA concentration was determined. To sequence the un-transfected STARR-seq plasmid library as baseline, twenty PCR reactions with 100 ng STARR-seq library were performed (98°C for 45s; followed by 6-15 cycles of 98°C for 15s, 65°C for 30s, 72°C for 45s) with the KAPA Hifi Hot Start Ready Mix and NEBNext Multiplex Oligos for Illumina (cat. no. E7335L).

### Next-generation sequencing

Next-generation sequencing was performed at the VBCF NGS Unit (<u>www.vbcf.ac.at</u>) on an Illumina HiSeq2500 platform. Genome-wide STARR-seq screens were sequenced as paired-end 50 cycle runs, focused BAC screens for signal-to-noise analysis as single-end 50 cycle runs, using the standard Illumina primer mix, STAP-seq screens were sequenced as paired-end 50 cycle runs with the TruSeq RP1 primer as a read 1 primer. All next-generation sequencing data is available at http://www.starklab.org/ and was deposited to GEO (accession number GSE100432).

### Measuring interferon-stimulated gene (ISG) expression levels

*Related to supplemental protocol: “qPCR ISG expression assay protocol”*

5×10^6^ HeLa-S3 or HCT-116 cells were electroporated with a focused STARR-seq library in three independent transfections (see section ‘transfection by electroporation’ for details). Mock electroporations without DNA were used as a negative control. 6h after transfection, cells were lysed using Qiashredder columns (Qiagen; cat. no. 79654) and total RNA was extracted from all cells using the RNeasy mini prep kit (Qiagen; cat. no. 74104), with beta-Mercaptoethanol supplemented RLT buffer. 1 μg of total RNA was treated with recombinant DNaseI (rDNaseI; Ambion, cat. no. AM1906) for 30 min at 37°C followed by the removal of rDNaseI using the DNase inactivation reagent provided in the kit (DNA-free DNA removal kit; Ambion; cat.no. AM1906). The DNaseI treated RNA was reverse transcribed using Invitrogen’s Superscript III (Invitrogen; cat. no. 18080044) and Oligo-dT_20_ primers (Invitrogen; cat. no. 18418020) (25° for 5 min, 50 °C for 50 min, 70 °C for 15 min). Gene expression levels were measured by qPCR on 2 μl diluted (1:5) cDNA using Go Tag SYBR Green qPCR Master Mix (Promega; cat. no. A6001) in a total volume of 20 μl with 0.5 μM gene-specific qPCR primers (95°C, 2 min; 95°C, 3s; 60°C, 30s; 40 cycles total). The sequences of all qPCR primers for ISGs and control genes (*ACTB, TUBB*) are listed in Table S1.

### qPCR based reporter assay on luciferase transcripts

*Related to supplemental protocol: “qPCR reporter assay Protocol”*

For each enhancer candidate, 5×10^6^ HeLa-S3 cells were electroporated with 18 μg of a firefly luciferase reporter plasmid and 2 μg of a Renilla luciferase control reporter plasmid (pRL-CMV, Promega; cat. no. E2261, see section ‘transfection by electroporation’ for details). After the 30 min recovery phase, C16 and BX795 inhibitors were added to the cells at 1 μM concentration (see section ‘inhibitor treatment’ for details). qPCR to quantify enhancer candidate driven firefly luciferase transcripts normalized to constitutively expressed Renilla luciferase transcripts was performed as described above, with the exception that we used Turbo DNase (Ambion; cat. no. AM1907) for cells transfected with reporter plasmids, to ensure removal of residual reporter plasmid DNA. qPCR primers for Firefly and Renilla luciferase transcripts are listed in Table S1.

## Computational Methods

### TSS determination of STARR-seq reporter transcripts using STAP-seq

To ensure equivalent sequencing-depth for all samples, all sequenced read pairs were randomly subsampled to 500,000 fragments using reservoir sampling^68^. After subsampling, for all sequenced read pairs, mate 1 was mapped to the respective STARR-seq plasmid backbone using bowtie^69^. For this, the 5’ most 8nt random barcode introduced during the RNA adapter ligation procedure (see Arnold et al.^8^ for details) was removed (but kept associated to its read pair) and the following 15nt that correspond to the 5’ end of the reporter mRNA were mapped uniquely allowing for 3 mismatches (bowtie option –v 3 -m 1 --best –strata). The resulting mapping locations were collapsed based on the random 8nt barcode, allowing for molecular counting of all reporter mRNA 5’ ends and therefore all detected initiation events. Finally, the sum of all initiation events per annotation region within the backbone (e.g. core promoter, origin of replication etc.) was calculated and collected in a tabular format. The percentage of core promoter initiation was calculated as a ratio between all initiation events within the ORI or the core promoter and all initiation events on the plasmid.

### Stratification of STARR-seq signal based on reporter initiation

Knowing the precise 5’ end of each reporter transcript and therefore the TSS within the plasmid based on mate 1, allowed us to stratify genomic fragments (determined by the 3’ mate of the paired-end sequencing) based on their origin within the plasmid (e.g. ORI vs. core-promoter). Mate 2 was then mapped to the human genome (bowtie options -v 3 -m 1 --best --strata). To create density tracks the resulting mapping location of mate 2 was extended by the average fragment length of the STARR-seq library used.

### STARR-seq data processing

STARR-seq single-and paired-end Illumina sequencing reads were mapped in fasta format to the human genome (hg19), only considering the regular chromosomes 1-22, and X using bowtie version 0.12.9 using bowtie options -v 3 -m 1 --best --strata^69^. Genome-wide screens without inhibitors and focused screens for signal-to-noise analysis were mapped as 50-mers, except the genome-wide screens with inhibitors that we mapped as 36-mers because of poor base call quality at the 3’ end of the read (with this adjustment, we achieved similar mapping rates). Only uniquely mapped reads with up to 3 mismatches and a maximal insert size of 2 kb for paired-end datasets were considered. Genome coverage bigwig files were generated using all reads with bedtools genomecov version 2.19.1^70^ and normalized to reads per million (r.p.m.). For both genome-wide STARR-seq and input libraries, reads from two biological replicates were combined.

### Peak calling

Enriched regions were shortlisted using all reads from the combined STARR-seq replicates versus input as described^7^ with a P-value cutoff of 1×10^−5^ and an enrichment cutoff 3 (enrichment over input, where input coverage was evaluated conservatively either locally at the peak summit or over a fixed input window of 20 kb surrounding the peak summit, whichever was higher) and peaks were then called with a corrected enrichment 4 (correction by the Wilson method as in Stark et al.^71^ to the conservative lower bound of a 99^th^ percentile confidence interval (z=3)). We discarded peaks for which a single fragment accounted for more than 50% of all fragments overlapping with the peak region. Peaks overlapping ENCODE blacklisted regions were also removed. Peaks were annotated as open if they had significant enrichments in Dnase I hypersensitivity datasets (binomial P-value < 0.05). This P-value was calculated over the entire STARR-seq peak window using the maximum DHS coverage (r.p.m.) over the median coverage in the input, a procedure that ensured that the summits of the respective datasets were evaluated and which yielded a FDR of 13.6% when applied to random regions.

### Comparisons of signal-to-noise ratios between different STARR-seq setups

To assess improvements in signal-to-noise upon changes in the STARR-seq setup or protocols (see main text), we needed a high-confidence set of functionally validated enhancers. In the absence of a large set of independently validated enhancers, we followed two strategies: we either evaluated STARR-seq signal-to-noise ratios on a small set of positive (n=4) and negative (n=6) regions according to luciferase assays. Alternatively, we defined a high-confidence subset of STARR-seq peaks in focused screens with support for endogenous enhancer activity based on DNase-seq and H3K27 acetylation. To this end, we shortlisted regions using all reads from focused STARR-seq screens versus input libraries from BACs (see section about input library cloning for details) as described above, but using a P-value cutoff of 1×10^−4^ and an enrichment cutoff 3. The peaks were then called with a corrected enrichment 4 (correction to the conservative lower bound of a 90.5^th^ percentile confidence interval (z=1.67)). A binomial P-value for DHS enrichment was calculated in a 20nt window around the peak summit for the median DHS coverage (r.p.m.) over the median coverage in the input of a window of 250. For H3K27ac the maximum coverage was identified in a 500 nt window around the peak summit. For this local maximum, a binomial P-value for H3K27ac enrichment was calculated as described for DHS. Peaks with an FDR adjusted P-value < 0.05 in both the DHS and H3K27ac were used as the predicted positive regions. DHS and H3K27ac data was downloaded from ENCODE (Table S2). The positive regions were then removed from the BAC regions and the remaining sequence was used as background. Positive regions for ORI with inhibitor, ORI without inhibitor and SCP1 were merged together and collapsed (bedtools merge –d 0) to make the analyses symmetrical for all screens and to prevent duplicate peaks within each cell type. The average coverage was calculated within each positive peak region then averaged across all positive peak regions, so all peaks would have equal weight. For the background regions, the coverage was summed and divided by the total number of nucleotides. This positive average was then divided by the background to give the final signal to noise value. For signal-to-noise bar graphs, the average score over all regions is displayed and error bars depict 75% confidence intervals. All P-values for signal-to-noise measurements between conditions were calculated using a two-tailed paired t test.

### Nearest TSS Gene Ontology

Peaks of interest were assigned to the nearest transcript TSS from the peak edges. Gene ontology analysis was done using topGO version 2.20.0^72^, which calculates significantly enriched terms using a Fisher’s exact test for these genes of interest over all genes as a background using Ensembl version 75 IDs^73^. Enrichments were calculated by dividing the significant number of genes in each term by the number of genes that one would expect by chance reported in the topGO output.

### Differential peak analysis

Peak calls for PKR/TBK1/IKK inhibited and non-inhibited genome-wide STARR-seq screens were combined and collapsed into one region if peaks overlapped by 85%. Differential peaks were called using a hypergeometric test; P-values were then adjusted with the Benjamini & Hochberg method (FDR).

### Motif analysis

Fasta files were generated for peaks regions of interest using bedtools getfasta^70^ for a 700bp window around the peak summits, considering only windows that were at least 50bp away from any transcript TSS (Ensembl version 75^73^). Motifs were called using MAST (Motif Alignment & Search Tool) from the MEME suite (Motif-based sequence analysis tools suite) version 4.8.1^74^ using options -hit_list -mt 0.00001. A motif was only counted once within each peak. Odds ratios and P-values were derived using a one-sided Fisher’s exact test. Down-regulated peaks (5-fold down-regulated, FDR adjusted P-value < 0.001) following treatment were compared to peaks that do not respond to treatment (fold change within +/-1.5 fold). The top and bottom 500 open peaks were compared to 9,613 random regions (see below). For motifs with FDR adjusted P-value < 1×10^−5^ the odds ratio is displayed colored by minus log_10_ transformed adjusted P-values.

### ChromHMM enrichments

The ChromHMM segmentations annotation was obtained from UCSC^36^. Each of the peak regions were overlapped with each of the ChromHMM candidate annotations terms (Enh, EnhW, ect.) using grep-overlap (http://compbio.mit.edu/pouyak/software; bed-tools intersect is also suitable), which reports the total positions each peak overlaps each annotation term. The overlap lengths were then summed for each annotation term and divided over the total length of overlapping all terms for either the all 9,613 high confidence peaks or all ranked peaks in non-overlapping bins of 500. Enrichments (binned or rank cutoff 9613) were then calculated by dividing these fractions by the fraction of the genome in each respective term either in the bin or the 9613 peaks. For binned plots, this was also done for the peak regions plus 50kb.

### Genomic distributions

Genomic annotation hg19 was downloaded from UCSC. Upstream was defined in this case as 2kb upstream of the first position of the first exon in a gene. Percentages were then calculated as mentioned above for ChromHMM above for all high confidence peaks.

### Heatmaps and meta plots

The average coverage was calculated in 50 bp non-overlapping windows for 40kb regions centered on STARR-seq peak summits using custom scripts in R. These regions were then sorted by the total occupancy in a 2kb window around the peak summit. Peaks were then separated by the presence of DHS over input with a binomial P-value < 0.05 for heatmaps and meta plots. Heatmaps were made from the sorted matrices using Java TreeView version 1.6.4^75^. Meta plots were constructed from coverage window averages using the colMeans function in R.

### Core promoter motifs in ORI

We searched for the occurrence of known core promoter motifs in the ORI by scanning the 620 bp long ORI sequence with position-weight matrices (PWMs) for 5 selected core promoter motifs conserved from fly to human^76,77^. PWMs for TATA-box, Initiator (INR), downstream promoter element (DPE) and E-box were obtained from Ohler et al.^77^ and PWM for TCT motif from Parry et al.^78.^ At every position along the ORI the sequence was scored against the respective PWM and the score was converted to the percentage of the maximal possible PWM score (perfect motif match). Strong motif matches (x 90%) were visualized along the beginning of the ORI sequence around the main initiation sites within the ORI.

### ENCODE RNA-seq processing

Raw fastq files were downloaded from the ENCODE RNA-seq dashboard. We only considered all cell lines for which polyA-selected total RNA was paired-end sequenced in at least 2 replicates. Protein coding transcript sequences (Gencode release 23^79^) were quantified using kallisto 0.43.0^80^ with sequence bias correction (--bias) and sample boot-strapping (-b 30). For each transcript, counts were normalized to sequencing depth and kept as counts per million (cpm), summarized to gene level and over replicates and then log_2_ transformed using EdgeR’s cpm function (prior.count = 2).

### ENCODE RNA-seq based clustering

Log_2_ transformed, gene-level counts (see above) of selected DNA and RNA sensor genes (see main text) were clustered using pheatmap 1.0.8^81^. The distance matrix was calculated using the maximum distance method. The hierarchical clustering was performed with complete linkage clustering.

### Normalized enrichment scores from i-cisTarget

Normalized enrichment scores (NES) for DNase-seq and ChIP-seq datasets were obtained from i-cisTarget^37^with default settings (minimum fraction of overlap of 0.4, ROC threshold of 0.005). We filtered all i-cisTarget results for DHS datasets (FAIRE & DNase-seq) for Encode DNase-seq datasets only and all ChIP-seq datasets for those performed in HeLa-S3 only. Any dataset with a NES cutoff > 3.5 (for DHS) or > 3.0 (for ChIP-seq) in any condition was considered for further analysis. To remove redundancy, only the maximum score for multiple scores from the same feature description was kept. The NES scores were then visualized using pheatmap (v1.0.8^81^).

### Repeat enrichments

Peaks containing repeat elements were identified by coordinate intersection with the UCSC RepeatMasker track for release GRCh37^44^ using bedtools. We required the annotated element to be entirely contained in the STARR-seq peak (-F 1.0), and at least 10% of the STARR-seq peak to overlap with the element (-f 0.1). Genomic background frequencies were calculated by coordinate intersection with 1×10^6^ randomly sampled genomic regions with the same size and chromosomal distribution as STARR-seq input fragments. Odds ratios and P-values for enrichments were calculated with a two-sided Fisher’s exact test using contingency tables of insertion frequencies. Adjusted P-values were calculated in R with the Benjamini & Hochberg method (FDR).

### Random control regions

We selected 9,613 random control regions from all possible fragments in the STARR-seq input library with a size distribution of 1000 to 1600 bp by reservoir sampling (sample version 1.0.2 https://travis-ci.org/alexpreynolds/sample). To match the peak size, we extended each region by 641 bp from the center of the fragment. For repeat enrichment analysis, we sampled 1×10^6^ regions to account for the low frequency of some elements and to avoid zero counts in the control set.

### qPCR analysis for ISG expression

Ct values for each target gene were normalized to the *ACTB* housekeeping gene using the delta Ct method described in Livak, & Schmittgen 2001^82^. Delta delta Ct values were calculated between electroporations with and without DNA and displayed in log_2_.

### qPCR analysis for reporter assay

Firefly luciferase Ct values for each candidate enhancer were normalized to Renilla firefly Ct values using the delta Ct method described in Livak, & Schmittgen 2001^82^. Delta delta Ct values were calculated between enhancer candidates and a negative control and displayed in log_2_. In the case of enhancer activity changes upon inhibitor treatment (Fig. 3H), delta delta Ct values were calculated between electroporations with and without inhibitor treatment.

